# Threshold trait architecture of Hsp90-buffered variation

**DOI:** 10.1101/016980

**Authors:** Charles C. Carey, Kristen Gorman, Becky Howsmon, Aaron K. Aragaki, Charles Kooperberg, Suzannah Rutherford

**Affiliations:** Division of Basic Sciences, Fred Hutchinson Cancer Research Center, Seattle, Washington 98109-1024, USA; Division of Public Health Sciences, Fred Hutchinson Cancer Research Center, Seattle, Washington 98109-1024, USA

## Abstract

Common genetic variants buffered by Hsp90 are candidates for human diseases of signaling such as cancer. Like cancer, morphological abnormalities buffered by Hsp90 are discrete threshold traits with a continuous underlying basis of liability determining their probability of occurrence. QTL and deletion maps for one of the most frequent Hsp90-dependent abnormalities in Drosophila, deformed eye (*dfe*), were replicated across three genetically related artificial selection lines using strategies dependent on proximity to the *dfe* threshold and the direction of genetic and environmental effects. Up to 17 *dfe* loci (QTL) linked by 7 interactions were detected based on the ability of small recombinant regions of an unaffected and completely homozygous control genotype to dominantly suppress or enhance *dfe* penetrance at its threshold in groups of isogenic recombinant flies, and over 20 deletions increased *dfe* penetrance from a low expected value in one or more line, identifying a complex network of genes responsible for the *dfe* phenotype. Replicated comparisons of these whole-genome mapping approaches identified several QTL regions narrowly defined by deletions and 4 candidate genes, with additional uncorrelated QTL and deletions highlighting differences between the approaches and the need for caution in attributing the effect of deletions directly to QTL genes.

---

Hsp90 chaperones the proper folding and interactions of a remarkable number of signal transducers in over 150 cell-cycle, development, transcriptional activation and chromatin remodeling pathways (http://www.picard.ch/downloads/downloads.htm) and conceals normally cryptic genetic variation in several organisms^1-3^. For example, reducing Hsp90 dosage by half in Drosophila results in the appearance of abnormal phenotypes in a few percent of flies in approximately 20% of genetic backgrounds in laboratory strains or wild populations ^4-6^. Different genetic backgrounds produce specific abnormalities, likely reflecting an abundance of normally-silent polymorphisms that quantitatively affect the strength of signaling through Hsp90-dependent pathways^1^. We recently provided evidence that Hsp90 specifically controls threshold traits, and suggest that the non-linear processes giving rise to thresholds may be a natural basis for the biological control of variation and canalization (the maintenance of invariant phenotypes despite genetic and environmental variation)^7^.

To better understand Hsp90 buffering and natural mechanisms of canalization, we have begun to dissect in detail the genetic and environmental bases of an Hsp90-buffered trait. Eye deformities are among the most common of a seemingly endless array of novel phenotypic abnormalities attributed to reduced Hsp90 function in Drosophila^6^. Here we dissect the genetic architecture of deformed eye (*dfe*) in replicate lines from a small founding population of a single *dfe* male and 3-4 unaffected sibling females from the same inbred cross. In the F_1_ generation, a handful of flies with eye deformities, and a similar number of unaffected flies, were selected and crossed amongst themselves to create separate high and low penetrance lineages. Subsequently, the high and low lines were each split into three lines, creating six highly related selection lines (HE1, HE2, HE3 and LE1, LE2, LE3). Each generation, penetrance (proportion affected flies) was scored and affected or unaffected flies were chosen as parents. A strong early selection response demonstrated the polygenic architecture and additive genetic basis of *dfe*^6^, while the Hsp90 mutation that first uncovered the trait was lost during selection through “genetic assimilation”^8^.

Genetic assimilation is easily understandable in the context of standard quantitative genetics by threshold trait models^9^, where a continuous underlying basis of genetic and environmental liability determines the expression of a small number of discrete phenotypic states dependant on thresholds^10^, such as result from non-linear developmental pathways and responses^11,12^. Environmental stresses, such as heat stress, or genetic disruptions, such as Hsp90 mutations, initially uncover abnormal phenotypes. Selection enriches the previously cryptic genetic determinants, increasing the number of individuals with genetic liabilities close to or surpassing expression thresholds, thereby increasing trait penetrance in the selected populations to the point they eventually express the traits independent of the original mutation or stress^1^. For example, the Hsp90 mutation that originally exposed the *dfe* trait is deleterious even to heterozygotes, and was quickly lost from the selection lines. Re-exposure of the selection lines to Hsp90 mutations further enhanced *dfe* penetrance, explicitly showing that Hsp90 is a genetic buffer of *dfe* polymorphisms^6^.

To determine whether the high lines had become fixed (homozygous) for *dfe* polymorphisms, at generation 47 HE1, HE2, and HE3 were each split into duplicate cultures, and selection was relaxed for 12 generations in one of each replicate. Penetrance remained high in either selected or unselected HE1 and HE3 lines, suggesting that *dfe* polymorphisms had become homozygous. By contrast, the average penetrance of HE2 decreased by about 30% under relaxed selection (*P*= 0.0008; Student’s T-test not shown), indicating that trait determinants were still segregating in that line. To limit the number of alleles segregating in each high line and to increase our mapping power, we reduced their genetic complexity by several rounds of intensive inbreeding and selection. Progeny of 10-12 single HE1, HE2, and HE3 females were scored for *dfe* penetrance, and the subline with the highest penetrance in each case was used to found the next generation of 10 or more isofemale lines. As shown in Table 1, after up to 5 rounds of selection, the coefficients of variation (right panel) decreased and penetrance (left panel) increased.

**TABLE 1.** Reduction of variation in *dfe* lines by isofemale selection.

| Line | Penetrance |  | Coefficient of Variation |  |
| --- | --- | --- | --- | --- |
|  | Before Selection | After Selection | Before Selection | After Selection |
| HE1 | 0.38 | 0.43 | 0.55 | 0.26 |
| HE2 | 0.32 | 0.73 | 0.61 | 0.14 |
| HE3 | 0.63 | 0.85 | 0.25 | 0.07 |

As selection progressed, the high lines developed slightly different morphologies. Because the lines differed from one another qualitatively in *dfe* phenotype and quantitatively in penetrance (fraction affected progeny), to determine the extent to which they shared *dfe* genes high and low lines were crossed *inter se* in all possible combinations. Penetrance was low among the hybrid progeny of crosses between any pair of low lines, and none of the low lines differed from any other in penetrance (Fig. 1; lower left quadrant). When high line HE1 females and males, or line HE2 males were crossed to any low line, their progeny had similarly low penetrance (≤10%) suggesting that the low lines lacked most recessive trait determinants. However, penetrance increased to 20-30% in the progeny of most low lines crossed to females from line HE2 or either males or females from line HE3, indicating that all low lines still carried some *dfe* polymorphisms that failed to complement the high lines, that HE2 and HE3 had higher breeding value for *dfe* than HE1, and that HE2 had a maternal effect. Further support for the maternal effect of line HE2 was seen in crosses amongst the high lines. Penetrance was uniformly high for any combination of high lines, (40-55% in most cases; upper right quadrant), but the maternal effect of HE2 in crosses with the other high lines was striking (Fig.1; compare the progeny of HE2 females (horizontally) with the progeny of HE2 males (vertically)). In crosses between HE2 females and any high line, from 63- 80% of the flies were affected due to a strong maternal effect (HE2 females X HE1 males > HE1 males X HE2 females, *P*_*two-sided*_=0.0357; HE2 females X HE3 males > HE3 females X HE2 males, *P*_*two-sided*_=0.0013; ANOVA not shown). These crosses differentiated the high lines in terms of breeding value (fraction affected progeny) and maternal effects, with HE2 ≥ HE3 > HE1. None of the variation for *dfe* segregated with the X chromosome, indicating that trait determinants were autosomal, and focusing our mapping efforts on the approximately 4/5^th^ of the Drosophila genome on the 2^nd^ and 3^rd^ chromosomes.

**Figure 1.**
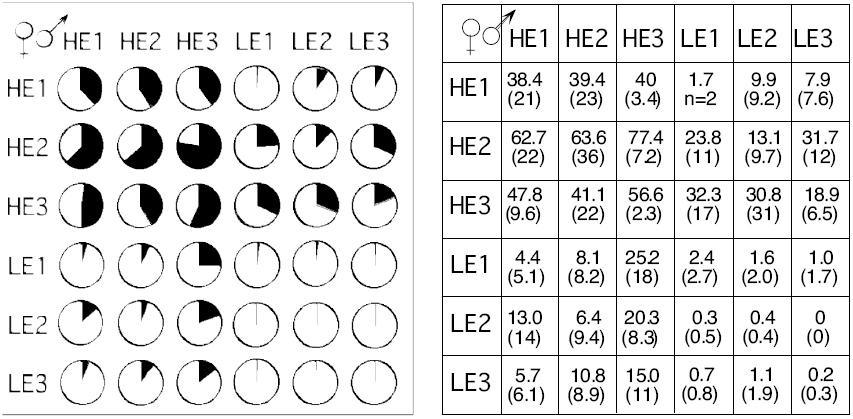
Heat map showing diversification of *dfe* lines in penetrance and maternal effect of HE2 line by complementation between control high (HE1, HE2, HE3) and low (LE1, LE2, LE3) *dfe* lines at generation 54 of selection. Maternal genotypes are shown horizontally; paternal genotypes, vertically.

A threshold model predicts that penetrance, the probability of *dfe* expression for a given genotype and environment, is determined by genetic and environmental liability and its proximity to the threshold. Therefore, near its threshold *dfe* alleles should contribute cumulatively and interchangeably to the trait while away from the threshold, only alleles with very strong effects on liability could affect phenotype. To test this idea, we created recombinant genotypes with increasing genetic liability. Through repeated backcrosses of a highly inbred and nearly isogenic wild-type line *Samarkand* (*Sam*) into each high line background, we created flies with on average increasing (but random) fractions of their genome from each high line. As with any unselected genotype, *Sam* flies never had deformed eyes, and in the F1 generation *dfe* penetrance was effectively 0% (just one *dfe* fly in 450 F_1_ flies scored), suggesting that many *dfe* alleles in the high lines are recessive to their *Sam* counterparts. Completely heterozygous and unaffected F_1_ females, hybrid for *Sam* and each mapping line, were crossed back to males from each respective high line, creating B_2_ flies heterozygous for a recombinant maternal chromosome made up on average of 50% *Sam* and 50% high line backgrounds. The B2 were again scored for penetrance and B2 females were backcrossed to their respective high line, generating B3 flies. With each generation of backcrossing, on average 50% of the *Sam* genome was lost by recombination and penetrance increased. As predicted, *dfe* responded to the increasing genetic (high line) variance or to increasing temperature in a similar non-linear (sigmoidal) response characteristic of threshold traits (Fig. 2). The inflection points of these curves identified a threshold for *dfe* expression: near 80% high line background at 25° C.—genetic liability, or near 26.5° C in the (100%) high line backgrounds—environmental liability). As shown in Figure 2, at the threshold penetrance increased sharply over a narrow range genetic or environmental liability, creating an almost binary phenotypic response.

**Figure 2.**
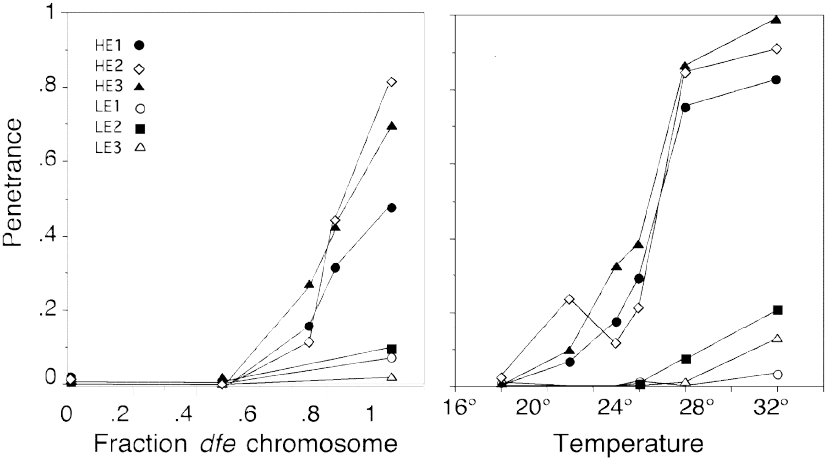
Genetic and environmental liability defining a *dfe* threshold. Replicate cultures of *dfe* flies from high (HE1-3) and low (LE1-3) lines grown at different temperatures (right), or grown at 25° C. with an increasing proportion of their genome from each *dfe* line in the *Sam* background (left). The graph on the right is reprinted with permission from Rutherford and Lindquist (1998).

We used the backcross lines closest to the *dfe* threshold, creating recombinant lines in which to QTL map the suppressive effects of WT *Sam* alleles on *dfe* penetrance. After 3 backcrosses an expected 7/8^th^ (87.5%) of each recombinant autosome was of high line origin. An additional 5 generations of free recombination was used to increase the genetic resolution of the mapping lines by increasing the number of meiotic recombination breakpoints^13^. In the penultimate generation, heterozygous recombinant females were crossed to males carrying an attached 2;3 balancer to extract unique pairs of recombinant chromosomes in single male flies. The recombinant, double balancer males were then backcrossed to high line females at 28.5° C., creating the recombinant isogenic lines that were scored for *dfe* penetrance and frozen for later genotyping. These were families of genetically identical recombinant flies from the same maternal and vial environments. Because of the backcross design, and the fact that *Sam* and each high line were very inbred and therefore nearly homozygous, only 2 genotypes were possible at any locus (*Sam*/high or high/high).

Figure 3 shows the distributions of *dfe* penetrance (probability of the *dfe* phenotype given each recombinant genotype) among 1,432 recombinant isogenic lines. The recombinant line phenotype distributions were greatly expanded from the narrow parental distributions of each *dfe* high line (Table 1), spanning the full range of possible penetrance from 0 to 1. Nearly 1/3 of the recombinant lines had 0 penetrance, identifying recombinant genotypes that strongly suppressed the *dfe* phenotype. Recombinant lines with penetrance higher than that of the parental lines suggested that some *Sam* alleles enhanced *dfe* penetrance in the high lines. The expected small size of the recombinant regions, breadth of the distributions and the overabundance of recombinant lines in the 0 penetrance class all suggested that the mapping design had successfully placed the recombinant genotypes in a range of liability near the trait threshold, allowing us to detect the effects of one or a few *dfe* QTL on penetrance.

**Figure 3.**
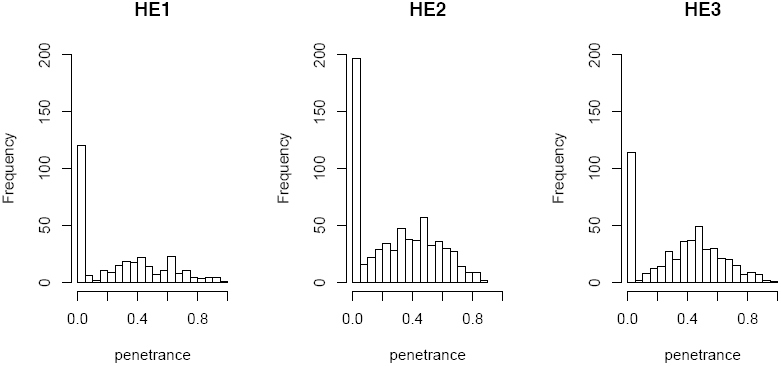
Penetrance of *dfe* distributed among 307 HE1, 664 HE2, and 461 HE3 recombinant isogenic lines.

To formally test this idea, a few dozen recombinant lines were selected from the 0 and highest penetrance classes for genotyping with markers of known molecular position across the 2^nd^ and 3^rd^ chromosomes, and at approximate densities of 5 cM (HE2) and 10 cM (HE1 and HE3). The genotypes of several recombinant lines indicated that they were heterozygous for just a single control (*Sam*) region. Assuming no undetected recombination, we reasoned that the recombinant regions in the 0 penetrance lines localized strong *dfe* allele(s) to intervals delimited by the positions of flanking markers. In every *dfe* line, low and high penetrance was most closely correlated with the genotype at genetic map position 2-69. When this position was homozygous for *Sam*, penetrance was high, and when that position was heterozygous, penetrance was almost always near 0, indicating strong dominant suppression by the *Sam* allele(s). We used the 2-69 marker to screen up to a few hundred randomly chosen recombinant lines from each *dfe* line. Comparison of the marker 2-69 genotypes and recombinant line phenotypes confirmed that a *Sam* allele(s) near 2-69 was a strong suppressor(s), with near 0 penetrance whenever the genotype was heterozygous. Since penetrance is bounded by 0 and 1, negative (suppressing) effects on penetrance beyond those of the *Sam* alleles at 2-69 were essentially un-measurable, regardless of the genotype at any other locus and whether or not additional suppressors of *dfe* were present. That is, because they already maximally suppressed penetrance, we reasoned that the *Sam* allele(s) near 2-69 were effectively epistatic to any other suppressing allele. To increase our ability to identify additional *dfe* QTL, we selected for full genotyping additional lines that were homozygous at 2-69 regardless of their penetrance.

Multiple interval mapping (MIM) was implemented by QTL Cartographer^14^ using a Bayesian Information Criteria (BIC) stopping rule. For each MIM model, a different BIC threshold was calculated based on the genome-wide 95% percentile of the maximum LOD score, estimated by permutation resampling^15^. Analysis of the selected and fully genotyped recombinant lines initially allowed us to detect 4 QTL in HE1 (139 recombinant lines), 3 QTL in HE2 (129 lines), and 4 QTL in HE3 (67 lines), confirming the presence of prominent QTL in the general vicinity of 2-69 in every *dfe* line; which in HE1 was centered at 2-70, in HE2 at 2-68, and in HE3 at 2-74. In terms of both their positive allelic effects on *dfe* penetrance, e.g. the difference in penetrance between high line and *Sam* genotypes, and their estimated effect on variance (PVE estimate) among the recombinant lines, the QTL near 2-69 had the largest effects on the *dfe* trait in any line. However, because the peak location of this QTL differed between the lines, we were uncertain whether it represented the same gene(s). We therefore initially had evidence for as few as 6 to as many as 11 *dfe* QTL in the 3 selection lines.

Given that the three *dfe* lines were expected to share many trait determinants, but had been handled separately and in parallel since they were first split at generation F3 of selection, they represented independent tests of *dfe* QTL positions and effects. Therefore, concordance of QTL in different *dfe* lines would increase our confidence in their significance. To confirm the possible correspondence between the *dfe* QTL and their interactions, we performed additional genotyping at four or five widely spaced markers in several hundred randomly selected recombinant lines, giving a total of 236 recombinant lines for the analysis of HE1, 450 for HE2, and 271 for HE3. All together, in this larger experiment QTL and their interactions accounted for 26 to 53% of the total variation in *dfe* penetrance. In HE1, the 4 previously detected QTL were again significant and located within 1 cM of their previous positions. We were able to detect an additional QTL on chromosome 3, giving a total of 5 QTL in HE1(Table 2). In HE2, the expanded analysis allowed us to detect three additional QTL, for a total of 6. This included the resolution of the major QTL previously centered at 2-68 into 2 peaks, a major QTL at 2-70 and another at 2-68. Additional QTL on the 2^nd^ and 3^rd^ chromosomes were found at 2-39 and 3-49, and the position of the QTL formerly at 3-68 was shifted to 3-65. However, for the HE3 line the situation was more complicated. Contrary to expectations, fewer QTL were detected when additional lines were genotyped and added to the analysis. The single QTL detected on the 3^rd^ chromosome disappeared with the addition of more recombinant lines, as did two QTL distal to the major peak on the 2^nd^ chromosome. However, the main peak in HE3 was more accurately localized, shifted from 2-74 to 2-70 in agreement with its localization in the other 2 *dfe* lines.

**TABLE 2.** QTL positions, interactions and effects, showing number of recombinant lines analyzed and approximate marker density (in parentheses).

| QTL <sup>a</sup> | HE1<br>139 (10 cM)<br>97 (40 cM) |  |  | HE2<br>129 (5 cM)<br>321 (40 cM) |  |  | HE3<br>67 (10 cM)<br>204 (40 cM) |  |  |
| --- | --- | --- | --- | --- | --- | --- | --- | --- | --- |
|  | Peak<br>location(s) | Effect <sup>b</sup> | PVE<br>estimate<sup>c</sup> | Peak<br>location(s) | Effect <sup>b</sup> | PVE<br>estimate<sup>c</sup> | Peak<br>location(s) | Effect <sup>b</sup> | PVE<br>estimate<sup>c</sup> |
| **49F;50D** |  |  |  | 2-67 | 0.21 | 16.6 |  |  |  |
| **50E;51C** | 2-70 | 0.38 | 53.2 | 2-70 | 0.16 | 17.8 | 2-70 | 0.24 | 23.1 |
| **90E;92E** | 3-67 | 0.17 | 0.9 | 3-65 | 0.14 | 7.4 |  |  |  |
| **27E;33B** |  |  |  | 2-39 | -0.1 | 1.8 | 2-38 | 0.16 | 9.3 |
| **61F;63B** |  |  |  | 3-2 | 0.08 | 2.2 |  |  |  |
| **71C;87A** |  |  |  | 3-49 | -0.08 | 0.7 |  |  |  |
| **93C;94D** | 3-73 | -0.19 | 4.4 |  |  |  |  |  |  |
| **43E;47A** | 2-59 | 0.11 | 9.1 |  |  |  |  |  |  |
| **21A;22D** | 2-0 | -0.10 | 0.5 |  |  |  |  |  |  |
| **(31F;32B)** |  |  |  |  |  |  | (2-43) | 0.31 | 22.3 |
| **(55B)** |  |  |  |  |  |  | (2-84) | 0.10 | 6.2 |
| **(57C)** |  |  |  |  |  |  | (2-93) | 0.24 | 18.7 |
| **(68F;69D)** |  |  |  |  |  |  | (3-38) | -0.70 | 1.3 |
| **E1** |  |  |  | 2-70x2-67 | 0.43 | 14.4 |  |  |  |
| **E2** | 2-70x3-67 | 0.27 | 16.5 | 2-70x3-65 | 0.2 | 10.8 |  |  |  |
| **E3** |  |  |  |  |  |  | 2-70x2-38 | 0.43 | 28.0 |
| **E4** | 2-70x2-0 | -0.21 | -3.8 |  |  |  |  |  |  |
| **E5** | 2-70x2-59 | 0.18 | 3.5 |  |  |  |  |  |  |

<sup>a</sup> QTL named according to the cytological locations of the most narrowly defined 1.5 LOD support interval or numbered epistatic interactions (E). QTL were arranged from highest to lowest effect with respect to the HE2 line followed by the other lines, and combined if they were within 1 cM of a central peak location. HE3 QTL in parenthesis, named for the cytological location of their peak position, were detected only in the reduced set of 67 selected fully genotyped lines or when lines homozygous at 2-70 were analyzed.
<sup>b</sup> Difference in penetrance between the high line and *Sam* genotypes at each position as calculated by QTL Cartographer.
<sup>c</sup> An estimate of the percent variance explained as % difference between the high line and *Sam* line at each position as provided by QTL Cartographer. The negative value for E4 interactions is an artifact of the method of calculation. Estimates of variance explained by the QTL model for each line, and the % variance explained by select QTLs or combinations of QTLs and reported in the text were calculated in R/QTL (Broman et.al., 2003). The QTL Cartographer and the R/QTL effect amount and % variance explained result from different methods and are not directly comparable.

For continuous and additive traits the addition of recombinant lines, whether sparsely genotyped or not, increases the statistical power to detect QTL^16^. However, for highly epistatic and threshold traits, which are buffered over a range of liabilities, the relationship between genotype and phenotype is complex, and can be completely uncorrelated over a range of liabilities away from the threshold. As there was some suggestion of additional QTL distal to the 2-70 QTL in the maps of all three lines (Fig.4), we reasoned that it was possible that the additional QTL detected in that region using the more limited numbers of extreme and selectively genotyped HE3 recombinants were in fact significant. Adding the randomly selected and sparsely genotyped lines to the analysis might have reduced the genotype-phenotype correlation in the data, obscuring the significance of previously identified QTL (e.g. Table 2, QTL in parentheses). Despite the inconsistent results of line HE3, overall the correspondence between the QTL detected in the three *dfe* lines was good. The higher resolution of the mapping data allowed us to infer that three QTL likely represent the same genes in multiple *dfe* lines. In summary, the straightforward MIM analyses had detected from 5 to 8 *dfe* QTL on the 2^nd^ chromosome (including the QTL distal to 2-70) and 4 or 5 *dfe* QTL on the 3^rd^ chromosome.

**Figure 4.**
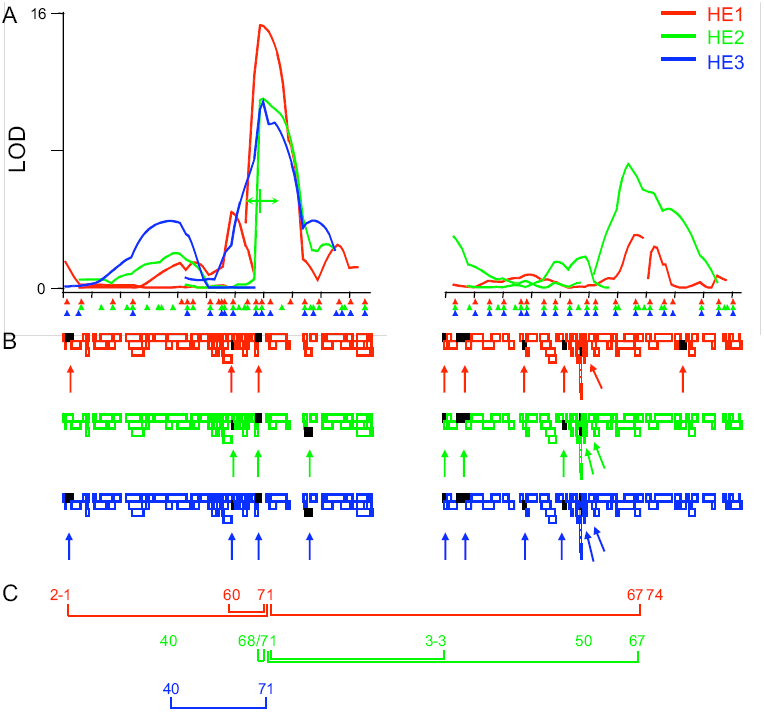
Correspondence of QTL and deletion maps across replicate *dfe* selections. A) Multiple interval mapping results from recombinants carrying *Sam* and HE1 (red), HE2 (green) or HE3 (blue) chromosomal intervals. Marker positions for each line are indicated by triangles. Chromosome 2 is left; 3, right. Tick marks are every 10 centiMorgans. Location equidistant between 2-68 and 2-71 is shown under HE2 curve in green to indicate that these two QTL were plotted by QTL Cartographer as a single peak. B) Deletions tested against the high lines are shown as empty colored rectangles in a staggered arrangement to maximize visibility of each deletion. Each rectangle indicates the portion of the genome deleted. Deletions with an effect on penetrance and described in Table 4 are shown as black rectangles (also indicated with arrows). C) Locations of the QTL are indicated by map locations, epistasis indicated by connective lines.

The major QTL at 2-70 was involved in every interaction (Fig. 4C), which together with its main effects accounted for up to 24% of the total variation for *dfe* penetrance. (By comparison, the QTL with the next largest effect on the model was at position 67 on chromosome 3 in the HE1 and HE2 lines, explaining approximately 5.3% of the effect in each case.) The epistasis of QTL 2-70 might indicate its position upstream of the other QTL in a genetic pathway. However, because of the threshold nature of *dfe,* we think it more likely that the epistasis of 2-70 simply results its large effect on liability and ability to bring other alleles near the trait threshold in that background, where even additive effects would cause the appearance of epistasis when they cumulatively impact *dfe* liability and penetrance (probability of the trait)^17^. To test this idea, we examined the data directly for evidence of alleles that were epistasic with the major QTL at 2-70 by repeating the MIM analysis on subsets of the previously analyzed selected and larger groups of more sparsely genotyped recombinant lines, eliminating the lines that were heterozygous for the strong suppressor at position 2-70 (thereby effectively shifting liability back toward the right and over the *dfe* threshold; Fig. 2). We reasoned that this would increase our power to detect suppressing *Sam* alleles with smaller effects on liability.

Interestingly, when we controlled epistasis by removing recombinant lines from the analysis that were heterozygous for the strong allele(s) at 2-70, several QTL changed. In addition to the expected disappearance of the 2-70 QTL in every line, several other peaks also disappeared, likely because of there dependence on the 2-70 allele to move the liability of recombinant lines toward the threshold where small changes in liability from additional QTL could affect *dfe* penetrance. The existence of an independent QTL centered at 2-68 in HE2 was re-confirmed by its existence independent of whether the 2-70 QTL was present or not. A broad QTL in HE3 was similarly subdivided into 2 QTL in this analysis, as the prominent peak at 2-38 disappeared and the originally more minor peak at 2-43 became significant. Indeed, the peak at 2-38 in line HE2 had a negative effect on the trait in line HE2, as opposed to the positive effect of the peak in HE3. Therefore, it seems probable that in line HE3 two QTL with opposite effects, one at 2-38 and one at 2-43, were confounded and masked by their interactions with the 2-70 QTL. If these inferences are correct, including all QTL found in line HE3 in either experiment, and the additional QTL detected by epistasis analysis directly, there were as many as 9 *dfe* QTL on the 2^nd^ chromosome and 5 on the 3^rd^ chromsome, for a total of 13 *dfe* QTL shown in Table 2. Similar analyses of the recombinant lines by composite interval mapping^18^ and standard permutation tests (CIM; not shown) identified and reconfirmed 10 of the same QTL found by multiple interval mapping (usually within 1 cM), including the detection of the two QTL distal to 2-70 in HE3, and additional QTL at 2-18 and 2-26, each identified by more than one analysis in one of three *dfe* lines, a QTL at 2-50 found in one of the analyses in one line, and a QTL at 3-38 found by several analyses in 2 of the *dfe* lines (HE2 and HE3). Combining all MIM and CIM analyses there is evidence for as many as 17 *dfe* QTL across the 3 lines.

Regardless of the exact number of *dfe* QTL, the QTL maps provided here show that the *dfe* trait is highly complex, particularly considering this genetic diversity represents a small sample of polymorphisms found in lines selected from a very small founding population of our *dfe* selection (1 male and 3-4 females). Notably, QTL with both positive effects (corresponding to suppressive *Sam* alleles) and negative effects on *dfe* (corresponding to enhancing *Sam* alleles) were present in about equal numbers and in all high line backgrounds. Given the unexpected abundance of suppressing alleles in the high lines (e.g. *Sam* alleles that enhanced *dfe* penetrance), we cannot rule out that some closely linked QTL with opposite effects were undetected by our recombination mapping approach. To independently identify and refine the locations of enhancers of *dfe* in each high line (mapped as *Sam* alleles that suppressed *dfe* penetrance) and to identify additional genes and pathways affecting *dfe* that were not polymorphic in the few flies that made up the founding population for the *dfe* selection, we tested each high line for complementation with the “Drosophila deletion set” in an approach similar to that used recently to estimate the number of bristle QTL in a mutation accumulation experiment in Drosophila^19^. At the time of testing, the autosomal complement of the deletion set included 155 ordered deletions covering approximately 85% of the 2^nd^ and 3^rd^ chromosomes (http://flybase.bio.indiana.edu/). Our deletion mapping approach took advantage of the fact that penetrance in the F1 progeny of crosses between high line and any unselected genotype was always less than 5%; increased penetrance would indicate the presence of a gene whose reduction shifts *dfe* liability, whether a *dfe* QTL or a gene involved in *dfe* pathways and processes.

We reasoned that deletions enhancing *dfe* penetrance would narrow the location of QTL with positive effects on the *dfe* trait, but we would not detect regions that suppress *dfe* penetrance (negative QTL effects), since penetrance was already near 0. Our initial criteria was increased penetrance in the progeny receiving the deletion, but not the balancer control chromosome. When we screened the deletion set close to the *dfe* threshold (at 29.5° C.) as many as 48 deficiencies enhanced *dfe* penetrance (≥ 5% affected) in the F1 progeny of females from one or more of the *dfe* high lines and males heterozygous for each deficiency (Table 3). Over 1/3 of these deletions corresponded to QTL regions broadly identified by recombination mapping. Because of the low resolution of QTL regions, and the high number of non-complementing deletions, many associations would occur by chance. To focus initially on those genes and deletions that were the strongest enhancers of *dfe* penetrance, we repeated the screen of the entire deletion set at lower temperatures (26.5° C. and 22° C.), effectively shifting *dfe* liability to the left, away from its threshold. As expected, this screen did not identify new regions, but defined a limited set of the deletions as “strong” enhancers of *dfe* penetrance (Table 4 and Fig. 4C), including Df(3L)M21 containing the single Drosophila Hsp90 gene, a previously established buffer of the trait^6^.

**TABLE 3.**
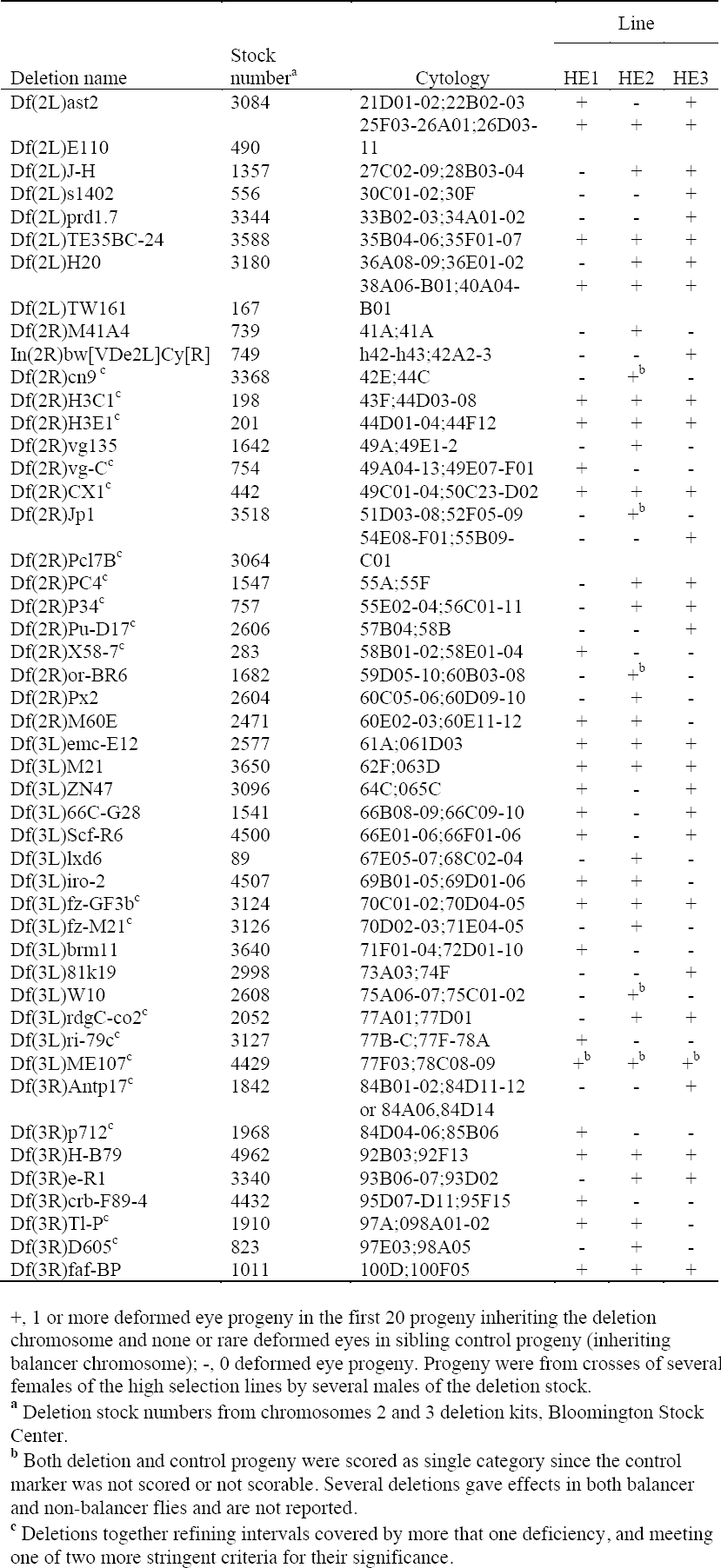
Weak (29.5° C.) enhancers of *dfe* penetrance.

The large number and high frequency of *dfe* genes likely segregate freely as neutral variants in many different genetic backgrounds. We believe this is because eye morphology is protected by a threshold, and depends on the strength of Hsp90 dependent signaling pathways and highly non-linear (and therefore canalizing) developmental responses. Because *dfe* genes are common, the effect of any particular deletion on *dfe* penetrance could be caused by (neutral) polymorphisms carried by the deletion chromosome, but unrelated to the deleted region. Therefore, our confidence in any particular region was greatly increased by the identification of additional overlapping non-complementing deletions. Several weakly interacting regions only identified at high temperature were defined by more than one deletion in the primary screen (Table 3; footnote c). To independently verify and confirm additional regions identified in the primary deletion screen, as well as to more narrowly identify regions controlling *dfe* penetrance, we fine-mapped deletion regions with independent sets of overlapping deletions. As shown in Figure 5, once a narrow interval was defined, all publicly available mutant alleles in the area were tested for the ability of individual genes to enhance *dfe* penetrance. This approach confirmed and restricted several regions defined by deletions to single or a few to a few-dozen candidate genes, revealing a rich and highly complex genetic architecture with a minimum of 14 regions and seven mutant alleles. Of 11 multiply confirmed regions or genes defined as strong enhancers of *dfe* penetrance (Table 4), seven lie directly within QTL identified in Table 2 by recombination mapping (from 0-3 cM from the most likely position of the QTL peaks). In short, the correspondence between the recombination and deletion mapping approaches employed here was good, but was not complete. These results underscore the importance of additional corroborative evidence, such as association studies correlating the trait with natural variants of the gene of interest in wild populations (as suggested by Mackay and colleagues20,21), before concluding that any gene or deleted region that fails to complement a quantitative trait corresponds to any trait QTL broadly spanning the same region.

**TABLE 4.**
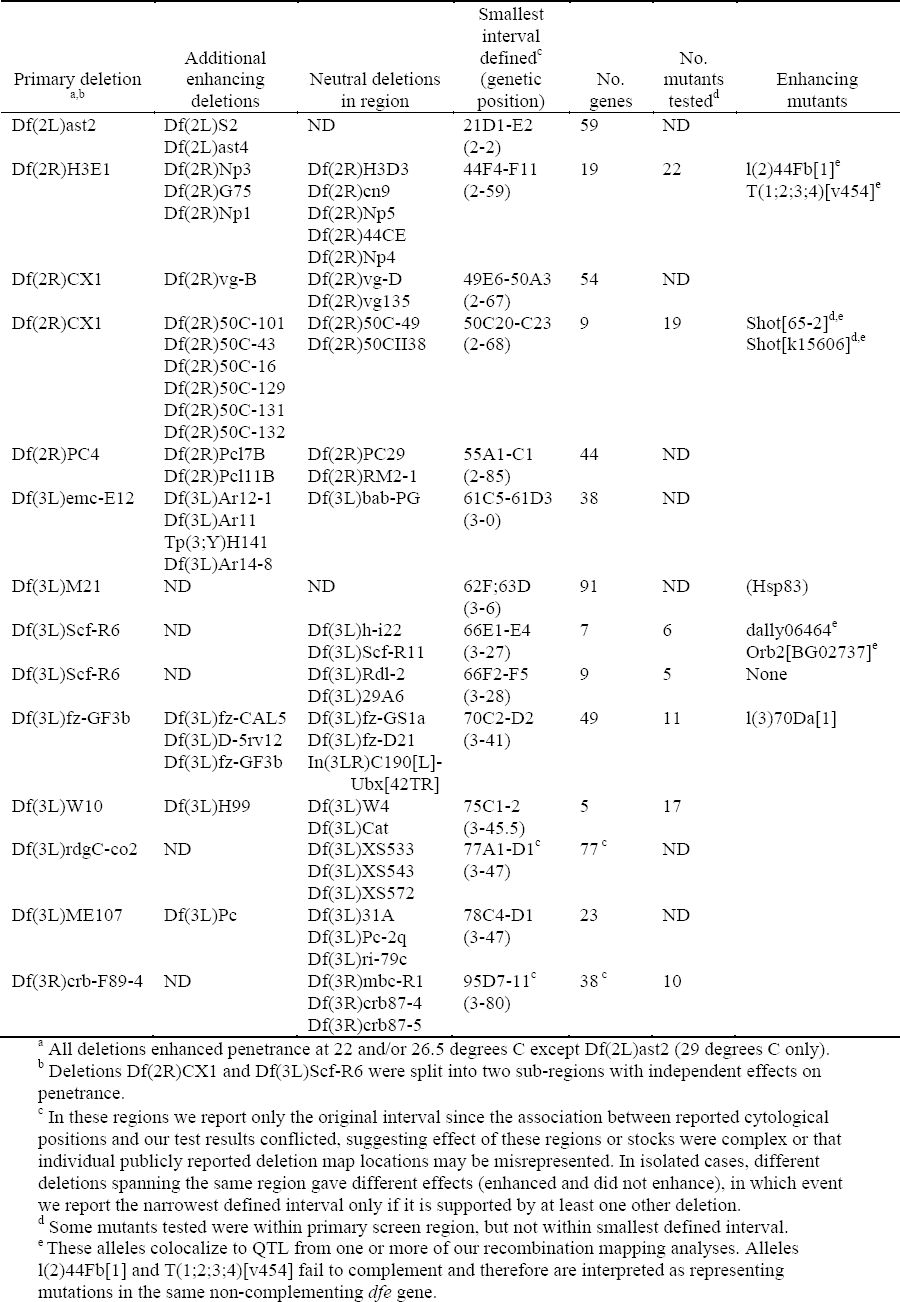
Strong (< 26.5° C.) and confirmed enhancers of *dfe* penetrance.

**Figure 5.**
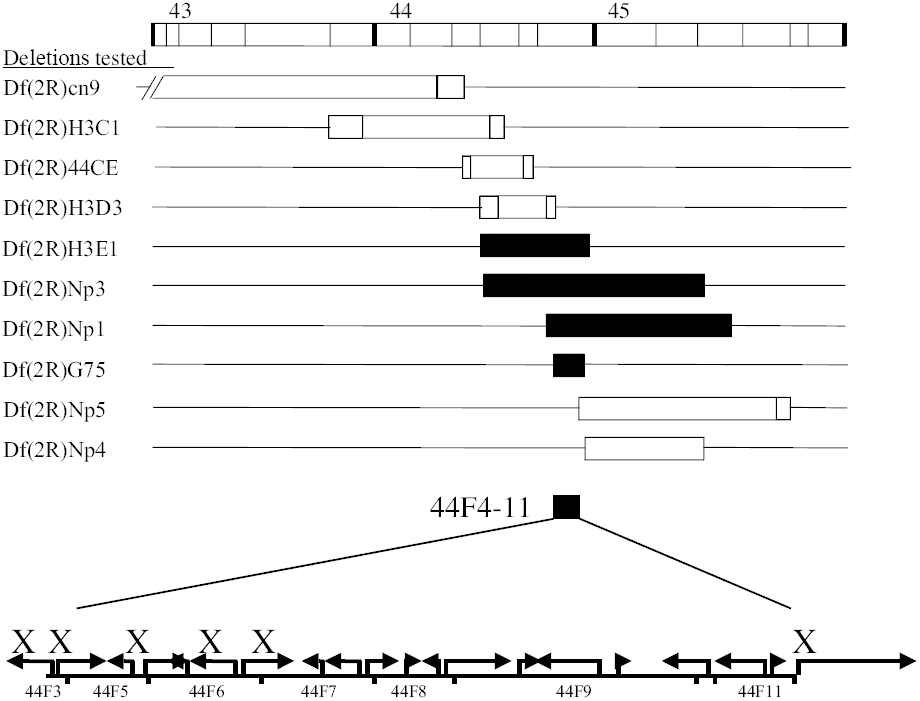
Dissection of region 44F to the level of a single penetrance enhancing mutant allele. Overlap between deletions with effects, delimited by neutral deletions defined cytological interval 44F4-11 as the narrowest subregion with an effect on penetrance. 22 mutant alleles for this 20 gene interval were tested for enhancement of penetrance. The mutant allele 1(2)44Fb[1] and the transposition T(1;2;3;4)[v454] enhanced penetrance (neither known to the sequence level and therefore not shown). The other 20 publicly available mutant alleles tested were neutral for penetrance. Six of the 20 neutral alleles are known to the sequence of sequenced gene level (X’s in figure indicating they did not enhance penetrance). Other mutants (including, l(2)44Fb and the transposition) map to this 20 gene cytological interval, but their parent gene is not yet known. The lethal allele l(2)44Fb[1] was further confirmed to have an abnormality in this interval as it was lethal in complementation tests with the enhancing deletions but not the neutral deletions (data not shown). The 12 genes to the right of this interval are prioritized candidate genes for attempts at transgenic phenotype rescue by germline transformation to identify the l(2)44Fb[1] gene or other dfe genes in this region. Cytological location is indicated at the top of the figure and below the blow-up region. Lines depict chromosomes; rectangles chromosomal deletions; stippling, uncertainty in cytological ends of deletion; black, enhancing deletions; white, neutral deletions.

## Discussion

Many developmental abnormalities and several human diseases have a probabilistic basis where no single genetic or environmental determinant is wholly predictive of phenotypic outcome, making the nature and identity of the polygenic variants for complex traits such as disease susceptibility remain largely unknown. This is particularly true for threshold traits, despite their importance for medicine and evolution. Recent work on threshold trait mapping has been theoretical, emphasizing model selection and statistical approaches^22,23^. By contrast, we took advantage here of the inherent nonlinearity of *dfe* penetrance and our knowledge of, and ability to control, genetic and environmental liability. Our experimental designs allowed us to sensitively detect genes that either enhanced (shifted liability to the right) or suppressed *dfe* penetrance (shifted liability to the left) in groups of genetically identical flies using standard QTL mapping models and complementation testing. While genetic and environmental effects are confounded by a simple low dimensional and linear concept of liability, our work here demonstrates that for organisms where genetic backgrounds and environments can be closely controlled the threshold trait model is a useful heuristic for experimental design. Although the threshold trait concept was introduced over 70 years ago by Sewall Wright to model morphological abnormalities in guinea pigs^10^, the application of threshold trait models to population genetics and evolution has been lacking. However, statistical inferences based on threshold trait models are now possible, and are beginning to be applied to evolutionary problems^24 in press^. Furthermore, recent progress in multidimensional molecular profiling may soon provide molecular proxies of disease liability that can be directly measured in human populations to predict disease states or therapeutic response. Threshold trait models based on these complex descriptors of liability will then be useful in guiding genetic mapping approaches whether or not the mechanistic details of the molecular markers for liability are well-understood.

The complex interactions between *dfe* alleles may be wholly attributable to the threshold nature of the *dfe* trait. The architecture of *dfe* is suggestive of a complex network of abundant, but normally neutral variation affecting eye shape in Drosophila. Once flies are near the *dfe* threshold, they are sensitized to additional, even small genetic, environmental and stochastic effects^7^. Indeed, *dfe* may be a modern manifestation of a polygenic system of genes controlling eye morphology in Drosophila called tumorous head (*tuh*), which arose spontaneously in a wild strain collected in Acahuizotla, Mexico in 1941. Since it’s discovery, both the original *tuh* line and another spontaneous occurrence of a similar trait were studied intensively over several years^25,26^. Strong selection for vision may lead to a particularly high degree of canalization and hidden genetic variation for eye traits. The threshold model predicts the rare expression of *dfe* type traits in nature and suggests how a reduction in Hsp90 buffering could expose multiple interchangeable variants that can be rapidly (if only artificially) selected independent of correlated fitness costs^7^. Indeed, the polymorphisms segregating in the *dfe* selection described here, and in the *tuh* lines described previously, are likely just the tip of the iceberg of genes that can affect eye morphology in Drosophila once buffering is reduced and liability is brought near eye morphology thresholds.

The statistical identification of deletions with additive effects identifying QTL responsible for normally variable bristle numbers and longevity was pioneered in a series of studies from the laboratory of Trudy Mackay^21^. The continuously variable bristle traits and life history traits they focus on almost certainly respond linearly to increasing genetic liability, and are largely unaffected by Hsp90^5,7^. By contrast, Hsp90-buffered traits are threshold traits (whether morphological traits such as *dfe* as shown here, or normally invariant bristle traits^27^). By definition, threshold traits are insensitive to changes in genetic effects over a wide range of liability and less amenable to standard statistical models. For example, of 48 deletions that affected *dfe* at high temperature, the effects of 3/4 of those (27) were masked at lower temperature (and liability). As noted previously by Mackay and colleagues for continuous traits, statistically significant effects of the deletion chromosome could be due to epistatic alleles carried by the chromosomes but unrelated to the deletion itself^20,21^. To identify genes and genomic intervals that are most likely to be involved in the *dfe* trait, here we applied a stringent criteria that deletions strongly interact (at low temperature) with *dfe* penetrance (Table 4), and that multiple independent deletion chromosomes define each region. For example, over 20 deletions, including 10 of the weakly interacting (high temperature) deletions listed in Table 3, met this 2^nd^ criteria. Fine-mapping of regions initially identified by the deletion set has given us five candidate genes whose mutations enhance *dfe*, and several narrowly defined intervals containing up to a few dozens of genes (as opposed to a typical QTL that contains ∼500 genes). Four of the candidate genes are located directly within trait QTL, and are currently being cloned in transgenic *dfe* lines using a threshold-based strategy similar to our recombination mapping approach, which will ultimately allow the unambiguous assignment of *dfe* QTL genes and alleles. The highly replicated and comprehensive comparison of QTL and deletion mapping approaches provided here suggests that while the sets of genes identified are overlapping, these approaches are not entirely congruent, and reinforces the need for additional tests to confirm the correspondence between polygenic variants identified by quantitative trait mapping and developmental mutations.

We gratefully acknowledge Trudy Mackay and the Summer Institute of Statistical Genetics at North Carolina State University in Raleigh, NC (S.R. and C.C.C.), funding from the 2002 Damon Runyon-Walter Winchell Cancer Research Foundation Scholar Award (to S.R.), NIH R01 GM068873 (to S.R.), NIH R01 CA74841 (to C.K.) and NIH T32 CA80416 (to C.C.C).

## MATERIALS AND METHODS

### *dfe* abnormalities

Flies were scored as normal or abnormal based on gross abnormalities in eye formation, ranging from the ectopic appearance of macrochetae, and subtle invasions of cuticle into the edge of otherwise normal eyes, to the nearly complete absence of eye facets. Segregation of characteristics particular to each subline was not easily quantifiable.

### Selection lines

High and low *dfe* lines were initiated with a cross between a single male with abnormal eyes to three sibling females from the same cross. Flies with or without deformed eyes were selected in single high and low lineages for four generations, when enough abnormal flies were available to split the selection into three independent high and low *dfe* selection lines (HE1, HE2 and HE3 and LE1, LE2, LE3). At generation 47, the high lines were split and carried under normal and relaxed selection for 12 generations.

### Recombinant isogenic lines

At generation 47 of the *dfe* selection, highly inbred derivatives of HE1, HE2 and HE3 were established through as many as 5 generations of isofemale selection. Twelve single mated females were isolated from each line and their progeny were scored for penetrance. The lines with the highest penetrance were selected, and the process was repeated, resulting in increased penetrance and decreased coefficients of variation (Table 1). Virgins of each line were then crossed to the highly inbred and nearly homozygous wild type strain *Samarkind I-236* (*Sam*; isogenized through over 236 generations of pair matings between sibling flies, a gift of T. F. C. Mackay). Completely heterozygous and unaffected F_1_ females, hybrid for *Sam* and each mapping line, were crossed back to males from each respective high line, creating F_2_ flies heterozygous for a recombinant maternal chromosome made up on average of 50% *Sam* and 50% high line backgrounds. This process was repeated two more times, generating flies with recombinant intervals averaging 12.5% of the *Sam* background in the context of each high line background. Eight backcross sublines were initiated within each line by crossing F_4_ males and females with either normal or abnormal eyes. The number of recombinant intervals and map distance were increased by allowing 5 more generations of free recombination during round robin matings between the sublines, to assure mixing of recombinant products. Since Drosophila meiotic recombination occurs only in females, the expected mean interval between recombination events was 20 cM, resulting from as few as 3, to as many as 7 female meioses. To isolate unique pairs of recombinant autosomes, recombinant females were crossed to an attached chromosome 2 and 3 balancer stock *yw; T(2;3)B3/Pin^88k^* (kindly supplied from the laboratory of S. Henikoff). Unique recombinant males were crossed to HE2 females at 28° C., creating families of recombinant isogenic flies. Heterozygous progeny lacking the balancer chromosomes were scored for the *dfe* penetrance (fraction abnormal over total) and stored at –80° C. for subsequent genotyping.

### DNA preparation

Drosophila genomic DNA was extracted from HE1, HE2, HE3 or *Sam* adult populations (the parental populations) using a modified version of the Wizard(®) Genomic DNA Purification Kit (http://www.promega.com/tbs/tm050/tm050.html), or from individual flies using a 50ul solution of 10mM Tris-Cl pH 8.0, 1mM EDTA, 25mM NaCl, 200ug/ul proteinase K^28^. Individual flies were fragmented in solution, incubated at 37°C for 1 hour, denatured at 95°C for 2 minutes to inactivate the proteinase K, and stored at -20°C.

### Genotyping and marker selection for chromosomes 2 and 3

Genomic DNA was amplified using PCR primers for sequence tagged sites (STS) designed by the Berkeley Drosophila Genome Project (BDGP) (http://www.fruitfly.org/SNP/). The STSs averaged 200bp in length and were chosen based on their specified genetic and cytological positions on chromosomes 2 and 3. Standard PCR methods, as described by Varian Chromatography Systems for use with AmpliTaq Gold (Applied Biosystems), were used for amplification.

To identify heterozygous (*Sam*/high line) versus homozygous (high/high) regions, we screened for STS markers that varied between *Sam* and each selection line background using denaturing high performance liquid chromatography (DHPLC) on DNA amplified from completely homozygous or heterozygous flies (ref). Denaturing high performance liquid chromatography (DHPLC) was used to determine the genotype (presence or absence of a single nucleotide polymorphism), of PCR amplified DNA sequences. Samples were run on either a Helix column (Varian Chromatography Systems) or a DNAsep column (Transgenomics, Inc.), both of which are capable of high-resolution DNA separations. The gradient conditions for each sample was identical, however, the temperatures differed for each STS based on their individual melting temperatures, determined using the Stanford Genome Technology Center website: **http://insertion.stanford.edu/melt.html**.

For genotyping we used sequence tagged sites (STS) that were generated and unambiguously placed on genetic and molecular maps through the Drosophila genome project. STS are of 350-400 bp sequences randomly positioned throughout the genome and defined by convergent oligonucleotide primers. Markers were chosen using the following three-step process: (1) Prescreen control DNA isolated adult flies with the genotypes *Sam*/*Sam*, Selection/Selection (e.g. HE1/HE1) and *Sam*/Selection line (e.g., *Sam*/HE1) to identify SNPs between *Sam* and each *dfe* line and confirm that marker STS were homozygous for *Sam*/*Sam* and Selection/Selection. (2) Screen 10 individual flies from each of the three control lines at each marker identified in (1) to verify that flies within the same recombinant inbred line were genetically identical. (3) The reproducibility of our genotyping was confirmed for many markers in duplicate among the first fully genotyped 138 lines of HE1 recombinants (41 markers, 128 lines of HE2 recombinants (59 markers), and 66 lines of HE3 recombinants (37 markers).

### QTL Mapping

Recombination map values of genes in immediate proximity to the STS and alternatively by translating cytolocation to the recombinant map using the cytological table (http://www.flybase.org/maps/) were used to place markers on the genetic map. Since recombinant intervals were generated using a backcrossing scheme, and recombinant lines were likewise the result of a backcross, we chose a first generation backcross design for one trait as an approximate and simplified model for mapping the trait. This should lead to conservative estimates of LOD scores in intervals due to the lower estimated recombination between markers in the backcross model, compared to the higher rate of recombination expected in our experimental set, or the use of backcross or recombinant inbred models based on additional 5 generations of backcrossing.

Multiple interval mapping analysis (MIM), was implemented in Unix using QTL cartographer (version 1.17) as described in Weber et. al.^29^, described briefly below:

1. An initial models is scanned with m QTL and t epistatic effects.
2. Scan the genome for the existance of an (m+1)th QTL.
3. Scan for the (t+1)th epistatic effect.
4. Reevaluate signficance of each QTL in the model
5. Optimize positions of QTL in the model
6. Return to step 2 and repeat process until model does not change.

Statistical significance was based on the Bayesian information criteria (BIC). Specifically,

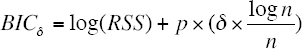, where RSS is the regression sum of squares and p is the number of terms in the model. Here δ was based on the 95^th^ percentile of the maximum LOD score, under the hypothesis that there are no QTLs. This 95^th^ percentile was estimated from permutation tests where the phenotypes are shuffled, keeping the genotype data intact^15^. Composite interval mapping (CIM; results not shown) was implemented using Windows QTL Cartographer v2.5 (http://statgen.ncsu.edu/qtlcart/WQTLCart.htm) using default parameters.

### Deletion Mapping

Virgin females from the HE1, HE2 and HE3 lines were crossed to males from deficiency stocks in the Drosophila Deficiency Kit (http://flybase.bio.indiana.edu/) covering approximately 85% of chromosomes 2 and 3. Each deficiency chromosome was heterozygous with a marked balancer chromosome. When possible, progeny of each cross containing the deficiency were scored separate from progeny containing the balancer chromosomes. When both deficiency and balancer chromosome classes had deformed eyes, a chi-square analysis was performed to determine if there was a significant difference between balancer and deficiency. Deficiencies having an effect on *dfe* penetrance were re-tested at 29° C with additional available overlapping deficiencies.

### Mutant analysis within deficiency defined regions

Narrow regions defined by fine mapping using overlapping deficiencies were subsequently investigated at the level of individual genes by crossing the females from the high lines to males from individual mutant stocks of genes in the region.

